# Single-cell transcriptome analysis of embryonic and adult endothelial cells allows to rank the hemogenic potential of post-natal endothelium

**DOI:** 10.1101/2021.06.16.447688

**Authors:** Artem Adamov, Yasmin Natalia Serina Secanechia, Christophe Lancrin

**Affiliations:** European Molecular Biology Laboratory, EMBL Rome - Epigenetics and Neurobiology Unit, via E. Ramarini 32, 00015 Monterotondo, Italy; Moscow Institute of Physics and Technology, Institutskii Per. 9, Moscow Region, Dolgoprudny 141700, Russia

## Abstract

Hematopoietic stem cells are crucial for the continuous production of blood cells during life. The transplantation of these cells is one of the most common treatments to cure patient suffering of blood diseases. However, the lack of suitable donors is a major limitation. One option to get hematopoietic stem cells matching perfectly a patient is cellular reprogramming. Hematopoietic stem cells emerge from endothelial cells in blood vessels during embryogenesis through the endothelial to hematopoietic transition. Here, we used single-cell transcriptomics analysis to compare embryonic and post-natal endothelial cells to investigate the potential of adult vasculature to be reprogrammed in hematopoietic stem cells. Although transcriptional similarities have been found between embryonic and adult endothelial cells, we found some key differences in term of transcription factors expression. There is a deficit of expression of *Runx1, Tal1, Lyl1* and *Cbfb* in adult endothelial cells compared to their embryonic counterparts. Using a combination of gene expression profiling and gene regulatory network analysis, we found that endothelial cells from the pancreas, brain, kidney and liver appear to be the most suitable targets for cellular reprogramming into hematopoietic stem cells. Overall, our work provides an important resource for the rational design of a reprogramming strategy for the generation of hematopoietic stem cells.

## Introduction

Every day, our body produces billions of blood cells to ensure the oxygenation of our tissues, to protect us from pathogens and to stop blood hemorrhage (Fliedner et al., 2002). This is possible because of the existence of hematopoietic stem cells (HSCs), rare cells residing in our bone marrow. They are characterized by the ability to self-renew and to produce any kind of blood cells. As we get older, our HSCs could slowly accumulate somatic mutations; some of which could lead to blood cancer (Marsilio et al., 2018). One strategy to treat this disease is to replace the entire hematopoietic system with a healthy one.

Every year, thousands of patients require hematopoietic stem cell transplantation to cure non-malignant diseases (e.g. thalassemia) and blood cancers such as acute lymphocytic leukemia (Passweg et al., 2019). These transplantations are life-saving but not every patient needing one can benefit from it because of the difficulty to find a matching donor. As a result, other methods to obtain blood cells matching a given patient have been explored. Cellular reprogramming is an approach where a cell from a person could readily be reprogrammed into the cell type of choice. The use of this approach would circumvent the need to find a compatible HSC donor. However, to successfully produce HSCs through reprogramming, it is crucial to understand how they are made in the body.

The generation of blood cells during embryogenesis is a very complex process involving the generation of many types of blood cells at different times and at different locations (reviewed in Medvinsky et al., 2011). HSCs are produced from endothelial cells, the building block of blood vessels, through a process called Endothelial to Hematopoietic transition (EHT) (Jaffredo et al., 1998; Zovein et al., 2008; Chen et al., 2009; Eilken et al., 2009; Lancrin et al., 2009; Boisset et al., 2010; Kissa et al., 2010). This transition occurs in the embryo at major haematopoietic sites such as the Aorta-Gonad-Mesonephros (AGM) region and the Yolk Sac (YS) (de Bruijn et al., 2000; Oatley et al., 2020; Shvartsman et al., 2019).

It has been assumed that the EHT only happens in a narrow time window during embryonic development and does not take place after birth. However, a recent study showed that the EHT is active in the bone marrow of chicken and mice after birth even though it eventually stops (Yvernogeau et al., 2019). This very surprising and intriguing discovery shows that the EHT could occur in non-embryonic conditions. Furthermore, murine adult endothelial cells from the lung were successfully converted into HSCs using a combination of transcription factors including Runx1 and Gfi1 and a cell line used as a niche in cell culture (Lis et al., 2017). This study was instrumental in showing that adult endothelial cells could undergo EHT if they were exposed to the right conditions.

In the past five years, the tremendous progress in single-cell RNA sequencing (sc-RNA-seq) has allowed the generation of several cell atlases such as the Tabula Muris (Tabula Muris Consortium, 2018). We used this atlas in comparison to embryonic AGM datasets to make the first direct comparison between the transcriptome of adult and embryonic endothelial cells. We ranked the potential of adult endothelial cells to undergo the EHT based on their gene expression patterns and gene regulatory networks.

Our work provides an important resource for the rational design of a reprogramming strategy for the generation of HSCs ex vivo. Additionally, it provides an example of how existing datasets can be explored to generate novel knowledge and contribute to the advancement of regenerative medicine.

## Results

### sc-RNA-seq analysis showed a partial similarity between embryonic and adult endothelial cells

To find how similar endothelial cells from the embryo and adult tissues were to each other, we compared the single cells from the AGM region between E9.5 and E11 coming from three different datasets called Embryo_dataset_1 (Hou et al., 2020), Embryo_dataset_2 (Vink et al., 2020) and Embryo_dataset_3 (Zhu et al., 2020) with the ones from Tabula Muris (Tabula Muris Consortium, 2018) (Supplementary Figure 1). This atlas is composed of twenty tissues but only twelve were found to contain endothelial cells after clustering analysis (Figure 1a). We confirmed that in each of the datasets there was a valid population of endothelial cells, well-separated from the other clusters. In the three AGM datasets, a large population of cells, expressing endothelial marker genes was identified. Of note, cells co-expressing endothelial and hematopoietic genes was found only in two out of three datasets (Hou et al., 2020 & Zhu et al., 2020). We reanalyzed the data from Embryo_dataset1 (Supplementary Figures 2 & 3) while we used the clusters identified by Zhu et al. for the Embryo_dataset3 (see methods section). Only one endothelial cluster was found in the Embryo_dataset2 (Vink et al., 2020). For the further integration with adult tissues, we picked the clusters related to the EHT.

**Figure 1:**
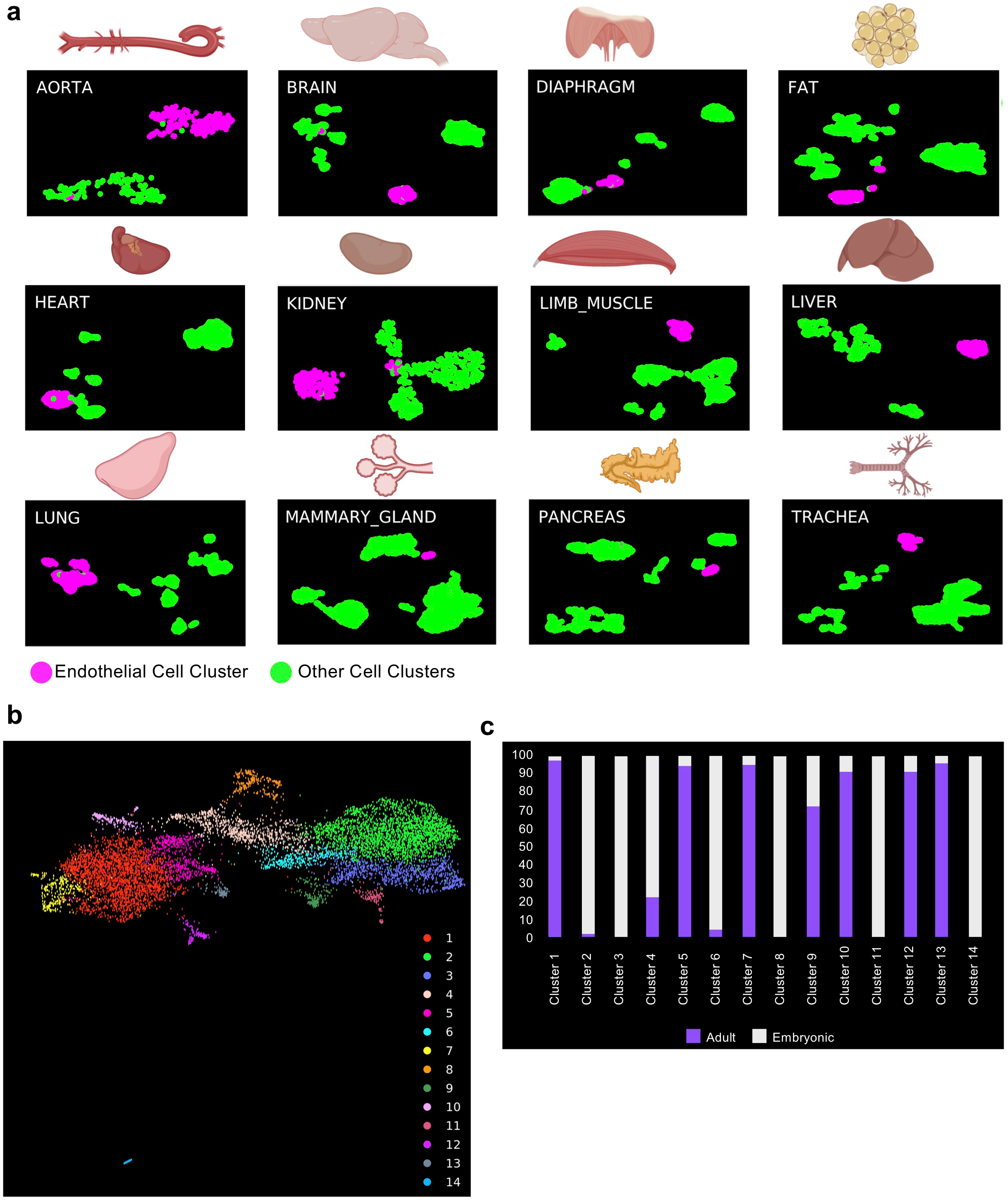
sc-RNA-seq analysis showed a partial similarity between embryonic and adult endothelial cells. **a)** UMAP plots showing the clustering analysis result for each of the indicated tissues; **b)** UMAP plot showing the clustering analysis result following integration of embryonic and adult tissues; **c)** Bar plot indicating the cellular composition of each cluster identified in b.

**Figure 2:**
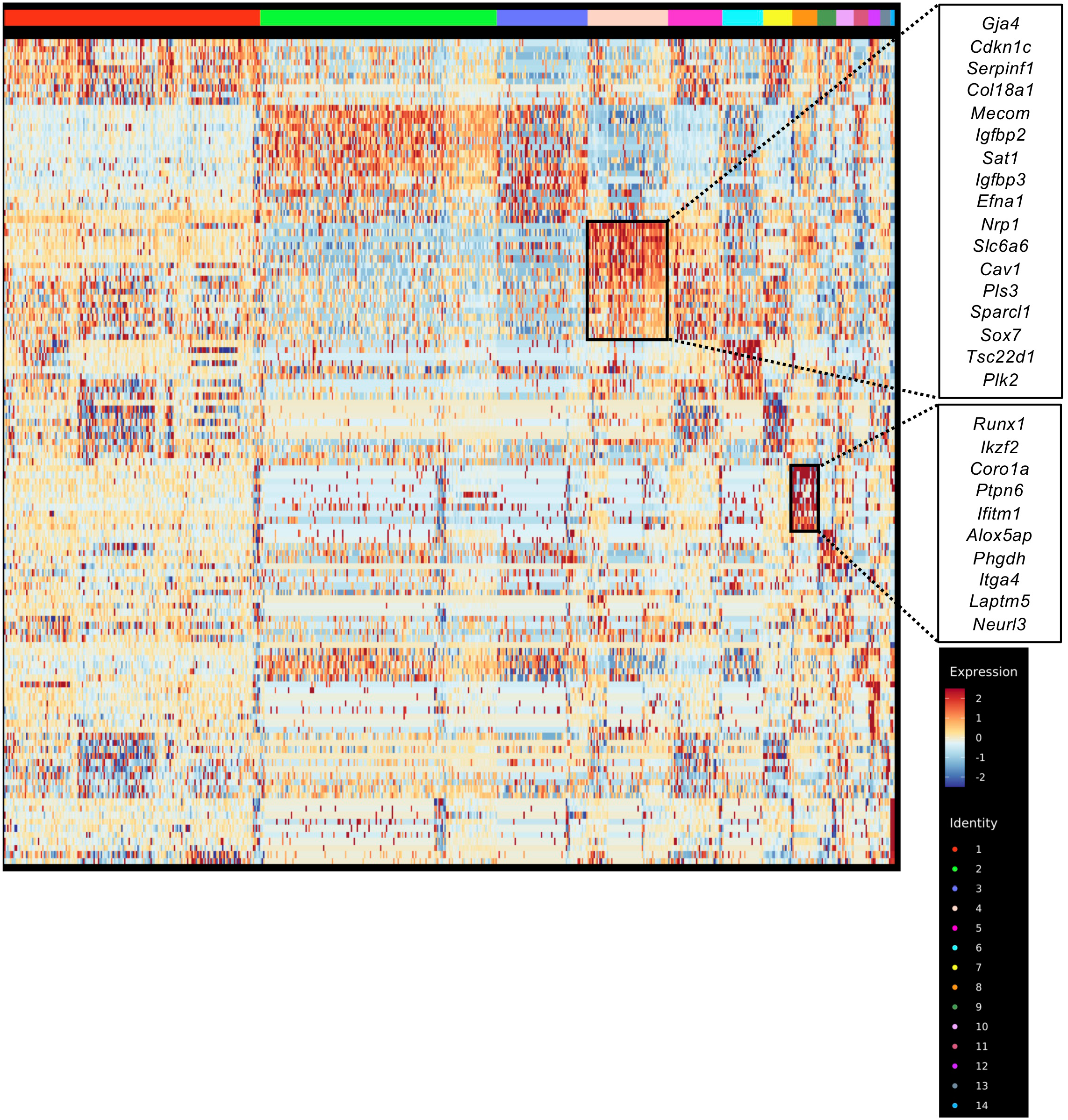
Embryonic Hemogenic endothelial cells cluster separately from all other endothelial cells. Heatmap of gene expression highlighting the top 10 marker genes of clusters 4 and 8 from Figure 1b.

The datasets were combined together using the data integration function of Seurat (Stuart et al., 2019). We then performed clustering analysis and identified fourteen distinct clusters from a total of 9541 cells (Figure 1b). We examined the relative composition of each cluster in relation to their origin (adult and embryonic). Out of fourteen clusters, eleven were composed of cells from adult and embryonic origin (Figure 1c). In contrasts, clusters 3, 8 and 14 were 100% composed of embryonic cells (Figure 1b & 1c).

An examination of the marker genes for each cluster helped us to determine their identity. Cluster 4 contains a mix of embryonic and adult endothelial cells with top 10 marker genes consistent with an arterial identity (Kalucka et al., 2020) (Figure 2). This is interesting because HSCs emerge from arteries during development (reviewed in Medvinsky et al., 2011). On the other hand, the cluster 8 which was exclusively composed of embryonic endothelial cells contained cells expressing hemogenic markers such as *Runx1* (Sroczynska et al., 2009), *Itga4* (Li et al., 2018) and *Neurl3* (Hou et al., 2020) (Figure 2). This supported the assumption that EHT only occurs in the embryonic tissues.

### Co-expression of *Erg, Fli1, Lmo2, Cbfb, Gata2, Tal1, Lyl1* and *Runx1* at the single-cell level is only detected in the mouse embryonic endothelium

Following our previous analyses, we did not find evidence of an EHT in the adult mice. However, we further asked how close adult endothelial cells could be to undergo the EHT process. Instead of basing ourselves on overall gene expression pattern, we specifically looked for the gene expression of key transcription factors (TF) crucial to the EHT process and hematopoiesis: *Cbfa2t3, Cbfb, Erg, Fli1, Gata1, Gata2, Ldb1, Lmo2, Lyl1, Runx1* and *Tal1*. In particular, the co-expression of *Cbfb, Erg, Fli1, Gata2, Lmo2, Lyl1, Runx1* and *Tal1* at the single-cell level was characteristic of the endothelial cells initiating the expression of hematopoietic genes (Bergiers et al., 2018). We therefore examined specifically how the genes coding these TFs were expressed in adult tissues in comparison to the embryonic ones.

Gene expression from endothelial populations was taken for expression and co-expression analysis in each tissue. As expected, the frequency of the TF gene expression was close to one hundred percent in embryonic endothelial cells (Figure 3a). That was especially striking for Embryo_dataset_1 and Embyo_dataset_2. Interestingly, the Embryo_dataset_3 showed lower frequency level compared to the other two datasets. This is likely connected to the 10X Genomics technology which cannot detect very effectively low expressed genes such as transcription factors (Wang et al., 2021).

**Figure 3:**
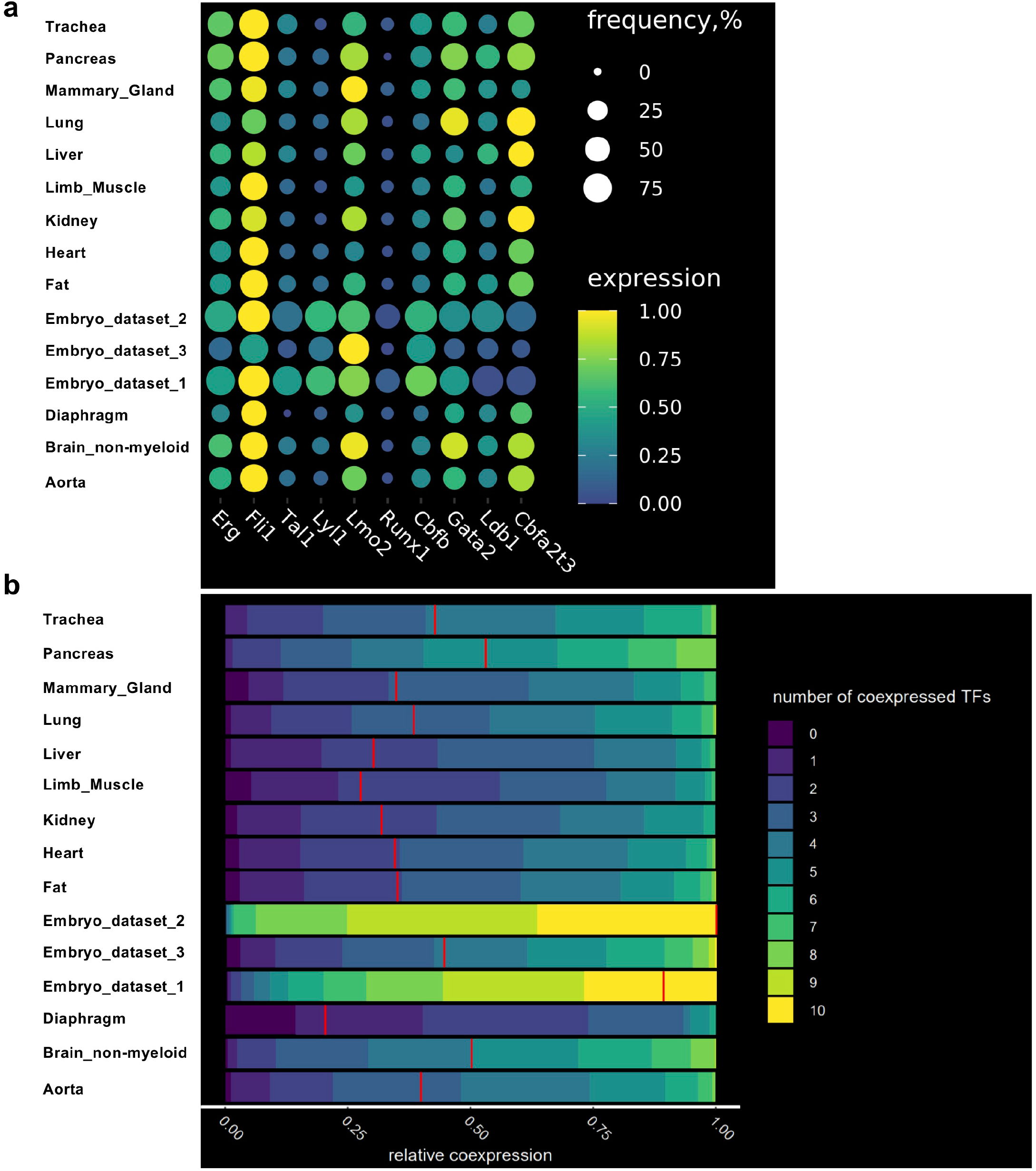
The frequency of cells co-expressing key transcription factors at the single-cell level is lower in adult endothelial cells compared to embryonic ones. **a)** Dot plots showing the frequency and level of expression of the indicated transcription factor coding genes in embryonic and adult endothelial cells; **b)** Heatmap showing the relative co-expression of the indicated transcription factor coding genes in embryonic and adult endothelial cells.

When we examined the endothelial cells from the adult tissues, we found that *Erg, Fli1, Lmo2, Cbfb, Gata2, Ldb1* and *Cbfa2t3* had moderate frequency of gene expression (above 25%) while *Tal1, Lyl1* and *Runx1* had low frequency (below 25%) (Figure 3a).

We next computed co-expression values of the TF gene expression in endothelial cells in embryonic and adult datasets. Consistent with the high frequency of TF gene expression (Figure 3a), embryonic endothelial cells from Embryo_dataset_1 and Embyo_dataset_2 revealed a high level of co-expression, more than 50% of the cells were expressing at least nine out of ten transcription factors at the same time (Figure 3b). The Embryo_dataset_3 showed much lower level of TF co-expression in line with the lower frequency of TF expression observed previously (Figure 3a).

Among the adult tissues, the highest levels of co-expression were observed in aorta, brain, lung, pancreas and trachea with about 50% of endothelial cells co-expressing five to seven TFs.

We questioned how the partial lack of gene expression of these transcription factors could affect their regulatory programs in endothelial cells. This problem led us to the inference of gene regulatory networks (GRN).

### Identification of putative target genes of the seed transcription factors in endothelial populations

To perform gene regulatory network analysis, we used the scTarNet R package that was developed for our previous study specifically focused on the mouse embryo (Bergiers et al., 2018). This method is based on choosing transcription factors as seeds for GRN analysis. We chose specifically the transcription factors that were examined above. Hereafter, they will be mentioned as seed TFs. A network was generated for each tissue where relationship between seed TFs and target genes was determined (Figure 4a). Of note, we only used the Embryo_dataset_1 along with the 12 adult tissues for this analysis. The Embryo_dataset_2 was not used because it did not capture the endothelial cells expressing blood genes. The Embryo_dataset_3 was excluded because of the lack of sensitivity of the 10X genomics technology (Wang at al., 2021).

**Figure 4:**
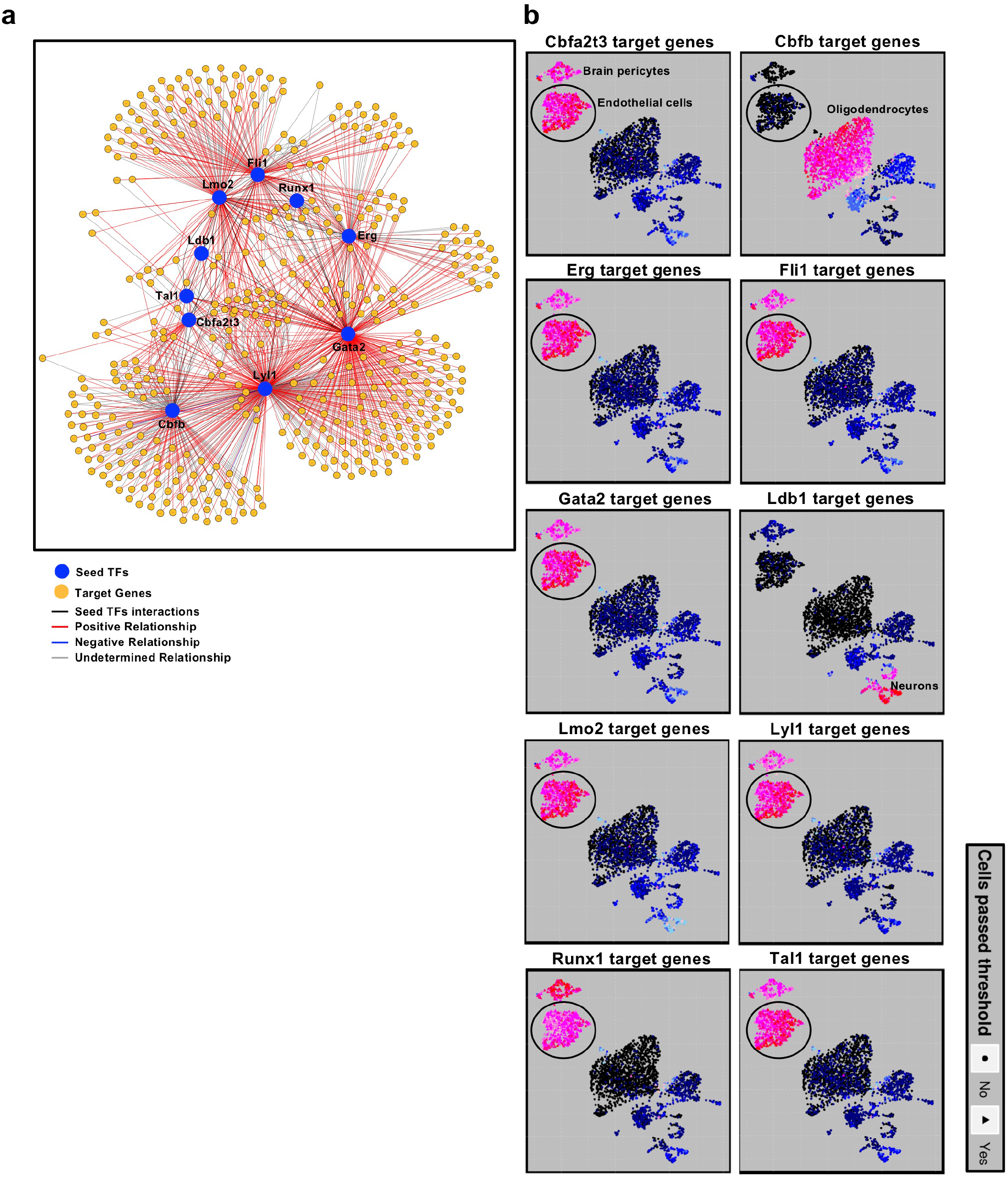
Gene regulatory network analysis identified target genes of seed transcription factors. **a)** Gene regulatory network generated by scTarNet. Blue circles indicate seed TFs, yellow circles indicate target genes. Red lines show positive relationship between seed TF and target genes; **b)** UMAP plot highlighting the clusters in which the target genes of a given seed TFs are expressed. Only positive relationships are highlighted. The ellipse shows the endothelial cell cluster. Colors represent AUC value – cells that passed AUC threshold colored in pink, others in blue.

To know in which cell clusters the seed TF – target gene interaction was happening, we specifically identified the cells in which the target genes were expressed. The results were highlighted in an UMAP plot. In Figure 4b, we show the results of this analysis for the brain. We did the same for each tissue and summarized the output in Figure 5. We found that seed TFs were positively associated with target genes, which were expressed in endothelial populations of almost every dataset, except diaphragm, fat and trachea. Aside from the Embryo_dataset_1, the highest grade of association between seed TFs and target genes were observed in brain, heart, kidney, liver and lung tissues.

**Figure 5:**
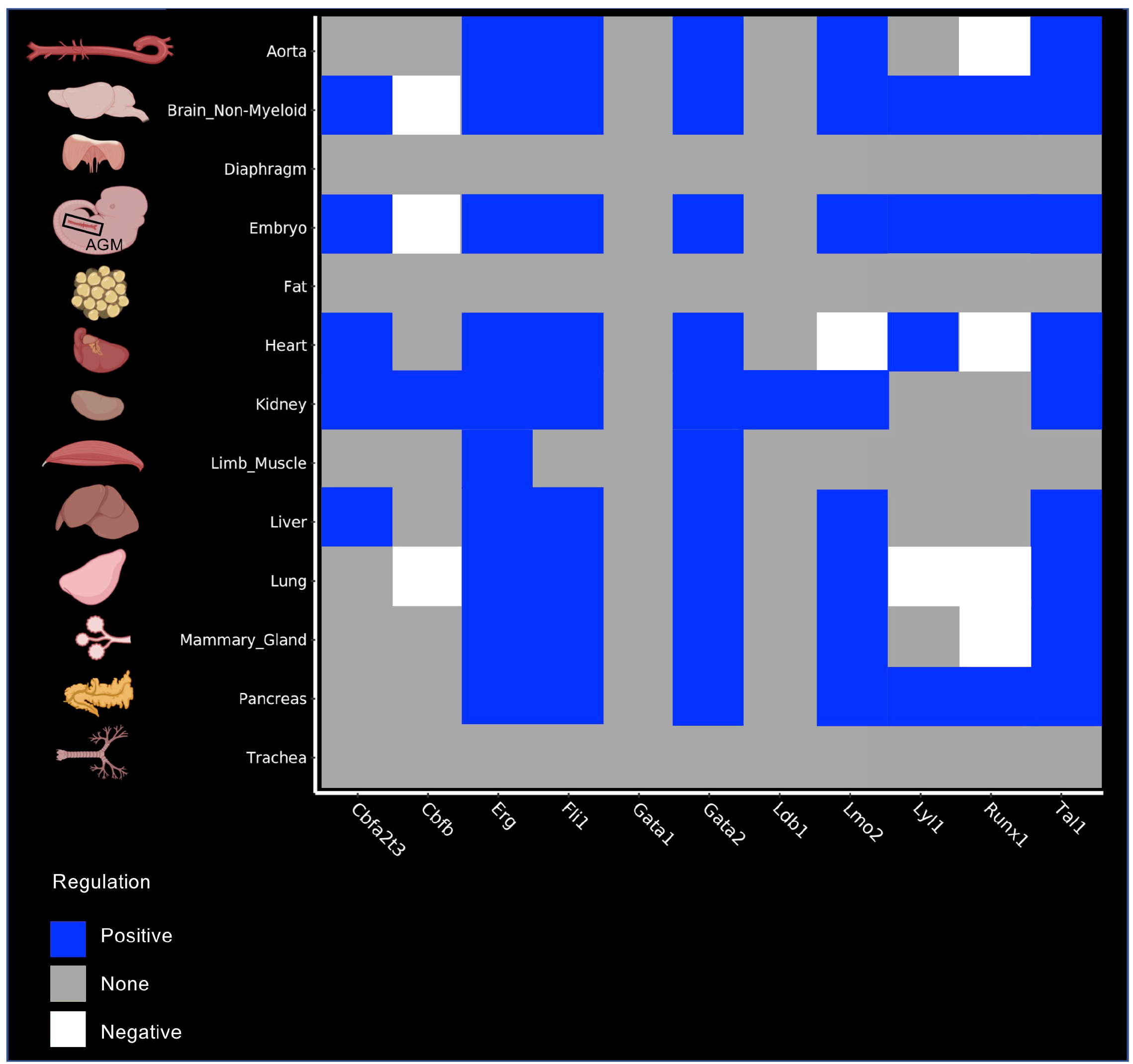
Identification of putative target genes of the seed transcription factors in each tissue. Heatmap summarizing the results of scTarNet for each indicated tissue. The colour code shows the type of regulation (positive, negative or none).

Although we identified a high level of association between TFs target genes in endothelial cells of adult tissues, it was not clear if there was any overlap between the groups of target genes connected to each seed TFs. Therefore, our next step was to perform a pairwise similarity analysis of these gene groups.

### Identification of common target genes between the different seed transcription factors

For each group of target genes associated with seed TFs obtained from the gene regulatory network analysis, we calculated pairwise overlapping genes between each group. Interestingly, the embryonic endothelial cells had the highest overlap of target genes between eight of out of eleven seed TFs (Figure 6). Among these eight seed TFs, Runx1 target genes were overlapping with the ones of Erg and Fli1 highlighting the existence of cells in transition between endothelial and hematopoietic cell fates. In contrast, this high-overlap was not seen in adult endothelial cells. However, the brain appeared to be the closest match to the embryo because we found a high overlap of target genes between seven out of eleven seed TFs (Figure 6). Of note, a clear difference was that Runx1 was not among these seven seed TFs. Following the brain, the heart, the liver and the mammary gland had an overlap of target genes for six seed TFs. The aorta, the kidney and the lung had one for five seed TFs. For these seven organs including the brain, Runx1 target genes did not have a high overlap with the other seed TFs (Figure 6).

**Figure 6:**
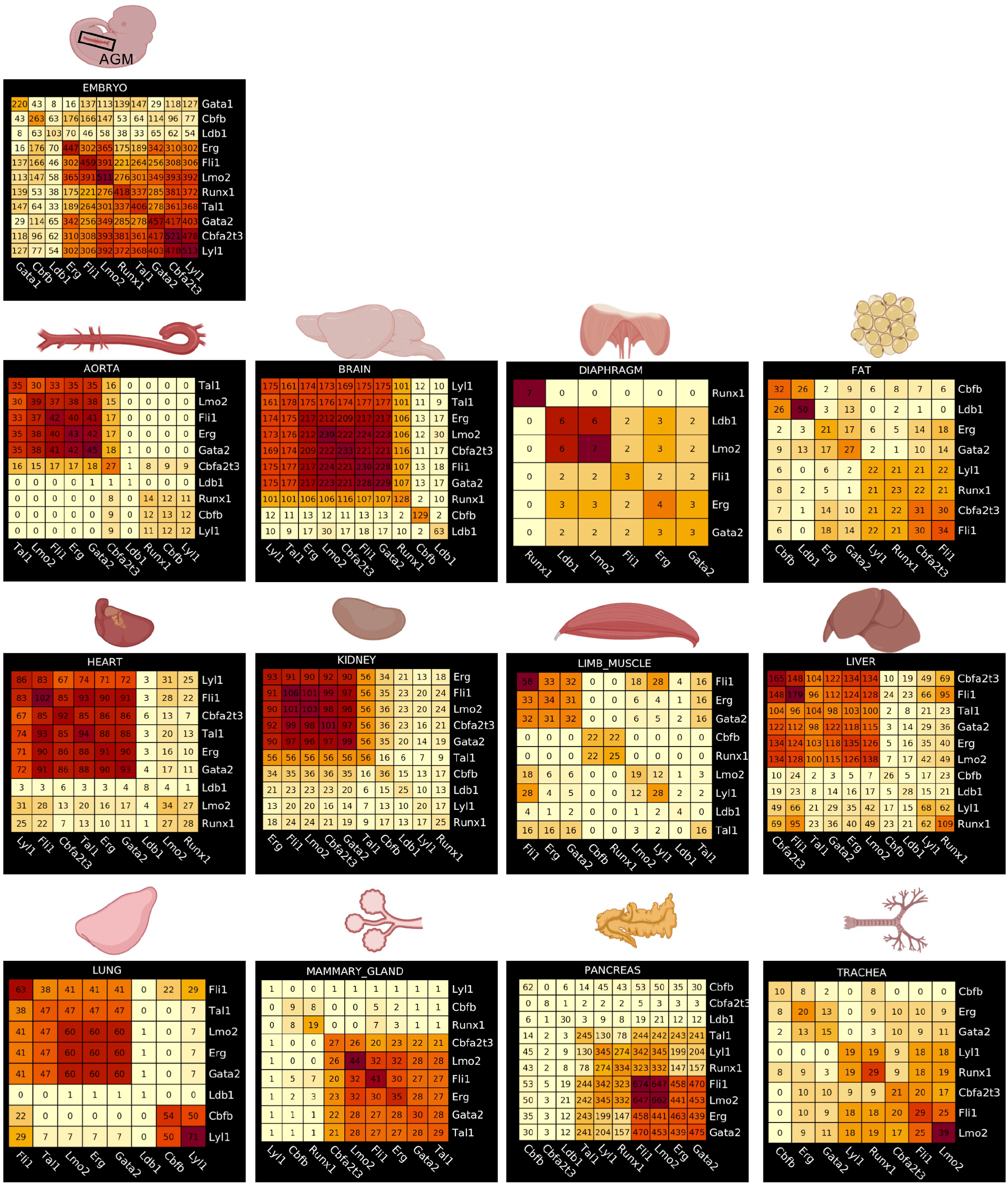
Identification of common target genes between the different seed transcription factors in each tissue. Heatmaps showing the result of a pairwise comparison of seed TF target genes in embryonic and adult tissues. Numbers in each heatmap correspond to the number of genes overlapping between two seed TFs.

The pancreas was presenting an interesting case. Indeed, we found a high overlap between four seed TFs (Fli1, Lmo2, Erg and Gata2). However, Runx1 target genes overlapped with about 50% of Fli1 target genes. When we examined more closely the results of the GRN analysis, we saw that this overlap of target genes between Fli1 and Runx1 was happening specifically in white blood cells (leukocytes cluster) and not in endothelial cells (Supplementary Figure 4).

In conclusion, our gene regulatory network analysis showed that many adult tissues had a quite high overlap of target genes between seed TFs. However, overlap with Runx1 target genes was rare.

### Runx1-specific clusters of adult endothelial populations have common marker genes

Runx1 is the main regulator of EHT (Lancrin et al., 2009; Sroczynska et al., 2009; Lie-A-Ling et al., 2018), that is why endothelial cells which were expressing this transcription factor were particularly interesting for additional investigation. Out of the twelve adult tissues, only the pancreas did not contain Runx1^+^ endothelial cells (Figure 3a). We performed clustering analysis of Runx1-expressing endothelial subpopulations versus the rest of endothelial cells in the eleven remaining tissues to identify the genes the most expressed in Runx1^+^ ECs (Figure 7a). We next compared each list to find the genes most frequently detected in at least four tissues (Supplementary Table 1). Only thirteen genes were found (Figure 7b). Some of them have been linked to hematopoiesis such as *Cd44, Notch2* and *Cd63* (Oatley et al., 2020) but most of them have not. There was no evidence of a hematopoietic cell fate in adult Runx1^+^ ECs. Of note, none of the key Runx1 targets in the EHT such as *Gfi1* and *Gfi1b* (Lancrin et al., 2012) were detected. These results suggest that *Runx1* gene expression is not sufficient to trigger the EHT process in adult endothelial cells.

**Figure 7:**
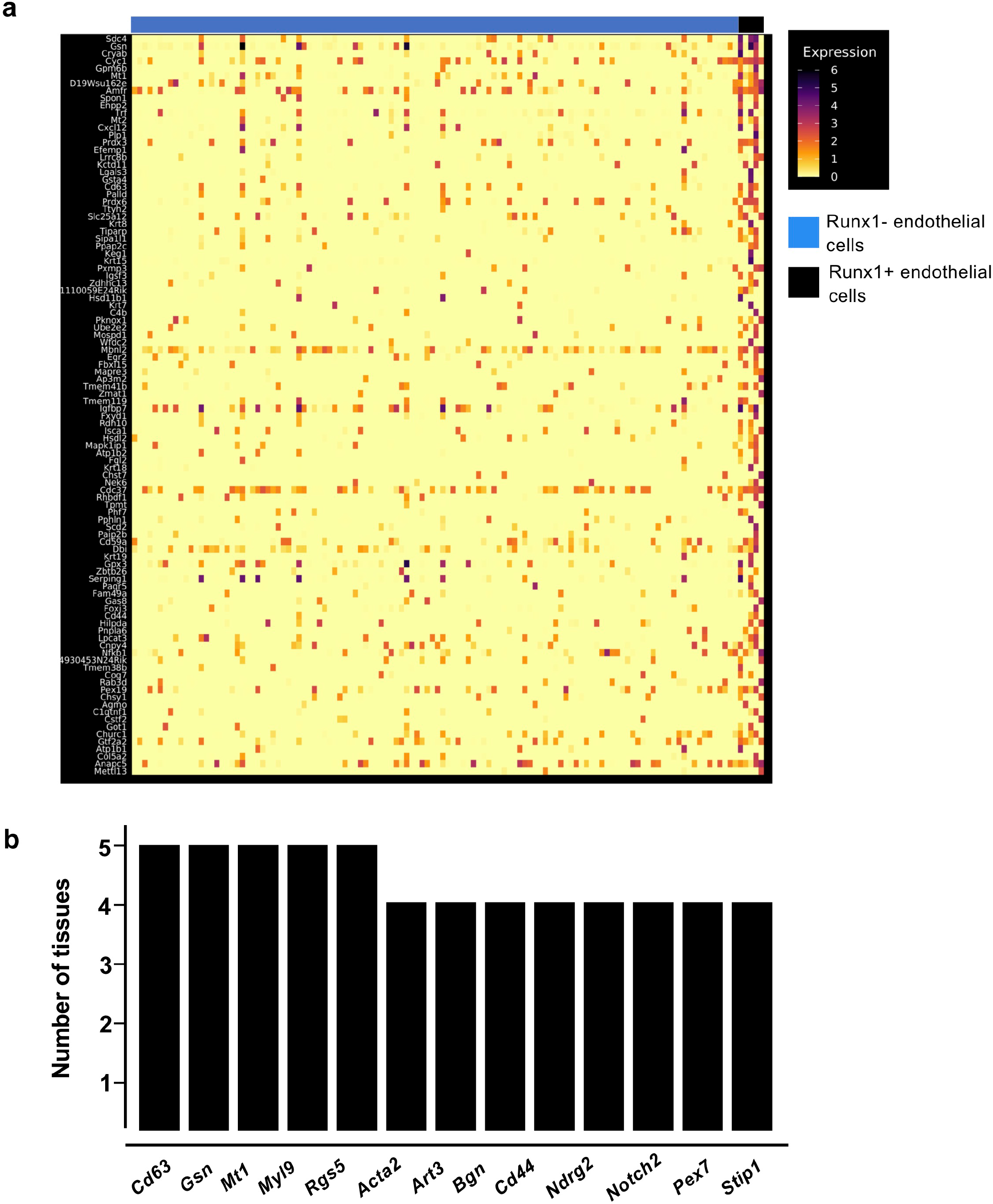
Runx1^+^ endothelial cells in adult tissues do not have a hematopoietic identity. **a)** Heatmap of gene expression highlighting the top marker genes of Runx1+ endothelial cells in the kidney; **b)** Bar plot showing the number of tissues in which the indicated genes are expressed in Runx1^+^ endothelial cells.

### The Pancreas, Brain, Kidney and Liver appeared to contain the endothelial cells the most suitable for reprogramming

Based on our gene regulatory network analysis, we have for the first time the possibility to rank the endothelial cells from adult tissues for their potential for EHT. We aggregated and summarized all the results that we had and came to the order of EHT-potentiality of adult mouse tissues based on: the expression of seed TFs in endothelial cells, co-expression of seed TFs and overlap between target genes of different seed TFs in endothelial cells (Figure 8).

**Figure 8:**
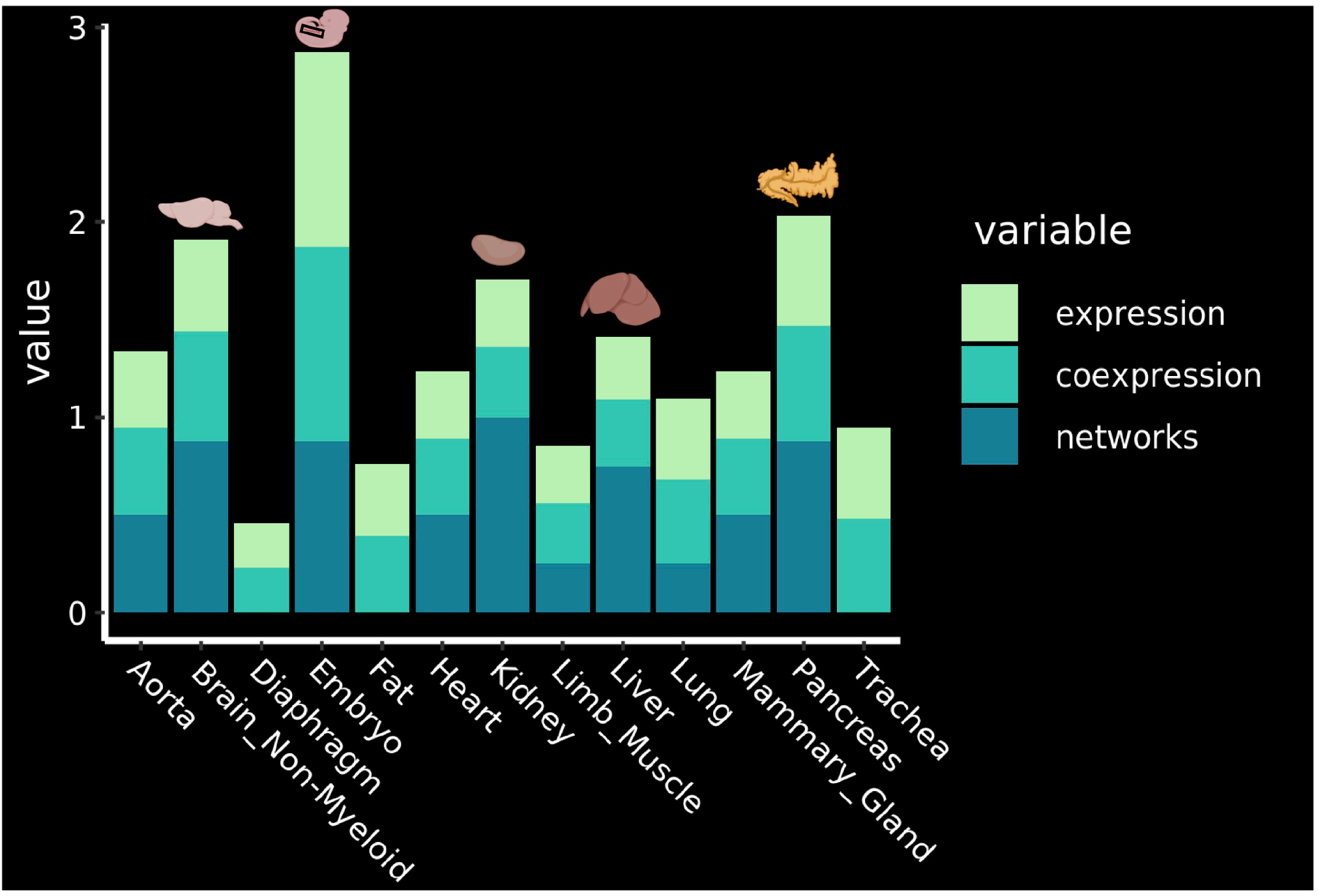
Identification of the most promising tissues for reprogramming. Ranking of the tissues according to the expression of the key transcription factors, their co-expression and the results of the network analysis. The top 4 organs are highlighted with their corresponding drawing. See methods for details about how the ranking was established.

The analysis showed that the four most promising tissues in which EHT might be triggered in the adult organism are the Pancreas, Brain Non-Myeloid, Kidney and Liver (Figure 8).

## Discussion

In this study, we investigated the process of endothelial-to-hematopoietic transition in the context of the adult mouse. We relied on the analysis of a number of high-quality single-cell transcriptomics atlases to address this question.

We first combined the datasets from the AGM, source of HSCs in the embryo, with the twelve tissues of the Tabula Muris where endothelial cells could be detected. Although we found that embryonic cells could cluster with a number of adult endothelial cells from different origins, endothelial cells with haemogenic characteristics could not be found in adult tissues. That reinforced the notion that the EHT is a transition occurring primarily at the embryonic stage.

We went on to determine which adult endothelial cells in the body were the closest to haemogenic endothelium. Instead of relying on overall transcriptional signature, we focused our study on key transcription factor coding genes *Cbfb, Erg, Fli1, Gata2, Lmo2, Lyl1, Runx1* and *Tal1*. These genes have been shown to be crucial to the EHT process. Besides, their simultaneous co-expression is enough to trigger the expression of blood and endothelial genes in vascular smooth muscles (Bergiers et al., 2018). Five of them, *Erg, Gata2, Lmo2, Runx1* and *Tal1*, were enough to reprogram fibroblast in hematopoietic progenitors (Batta et al., 2014). Their expression was therefore assessed in adult endothelial cells compared to the embryonic ones. As expected, none of them expressed all eight genes at the single cell level. Nonetheless, we noted differences between adult tissues. The aorta, brain, lung, pancreas and trachea displayed the highest level of co-expression with about 50% of endothelial cells co-expressing five to seven TFs.

There has been evidence that these TFs could work together as a complex helping to initiate the formation of blood cells from endothelium (Wilson et al., 2010 & Bergiers et al., 2018). In order to assess this possibility in the adult endothelial cells, we performed GRN analysis. The results from the embryonic endothelium supported indeed co-regulation of similar genes by these transcription factors. We also found extensive overlaps between the putative target genes of these TFs in the different tissues. Interestingly, the Brain ECs appeared to be closest to embryonic cells in that regard. Of note, the master regulator of EHT, *Runx1*, was found expressed at a low frequency in eleven of the twelve organs. However, none of the Runx1+ ECs had a hematopoietic identity suggesting that Runx1 alone was not enough for triggering the hematopoietic program in endothelial cells. Another clear difference with the embryonic stage is the relatively low expression of *Cbfb*, a key partner of Runx1. It was usually considered to have a ubiquitous expression in the embryo but in the adult it was not the case. This might explain why Runx1 is not triggering the hematopoietic program in post-natal vasculature. An additional key player is *Tal1*. Its role in the embryonic endothelium is to block alternative cell fates but it is also important for hematopoietic gene expression. We also noticed a much weaker expression of this gene compared to the embryonic stage. This could be another contributing factor to the lack of EHT at the adult stage. This is supported by a study on *Tal1* that we are currently performing in the group (Serina Secanechia et al, 2021).

The reprogramming of fibroblast or iPSC into HSCs has been very challenging. Some successful efforts have been reported recently and made it obvious that an endothelial stage is a prerequisite to the formation of HSCs *in vitro* (reviewed in Blaser et al., 2018). Murine lung ECs have been successful reprogrammed into HSCs but it was unclear if they were the best for such reprogramming (Lis et al., 2017). Our work offers a very valuable insight into this question, identifying which endothelial cells may be more suitable for reprogramming. This resource could be the stepping-stone for attempting in vivo reprogramming by targeting the most promising endothelial cells in the adult organism.

## Methods

### Datasets used in the study

For this project, we used four freely available sc-RNA-seq datasets of murine cells. The Embryo_dataset_1 (Hou et al., 2020; GSE139389) corresponds to embryonic tissues, as it contains 1432 single cells from 29 mouse embryos captured with high-precision single-cell transcriptomics. This dataset covers relevant EC populations at continuous developmental stages (E8-E11) which corresponds to the process of EHT. Single cell libraries were sequenced on Illumina HiSeq 4000 platform in 150□bp pair-ended manner. UMI-based scRNA-seq method was used to measure the gene expression profiles within individual cells. Cells expressing endothelial markers were used for data integration with adult endothelial cells.

The Embryo_dataset_2 (Vink et al., 2020; GSE143637) is sc-RNA-seq of AGM cells processed with the Smart-Seq2 protocol. The Endothelial cell cluster was used for data integration. The Embryo_dataset_3 (Zhu et al., 2020; GSE137116) is composed of single cell RNA-Seq profiles of cells involved in EHT from mouse embryos at embryonic day 9.5, 10.5, 11.5. Only the largest data subset called ‘E10.5 E+HE+IAC’ was used in our study. It corresponds to the clusters called ‘Conflux E’, ‘Endo’, ‘HE’, ‘IAC’ and ‘pre-HE’. Cells from the developmental trajectory of pre-hematopoietic stem cell formation was processed for library preparation using the 10x Genomics Chromium Single Cell 3′ Reagent Kit v2.

The last dataset was Tabula Muris that is an atlas of cell types from the mouse Mus musculus which comprises single-cell transcriptomic data from 100,605 cells isolated from 20 organs from three female and four male mice (Tabula Muris Consortium, 2018). We used only the part of this dataset which has cells sorted with FACS and libraries prepared with Smart-seq2 protocol. This protocol is a full-length transcriptomic technique that allows for more accurate analysis of the expression of transcription factors in comparison to droplet sc-RNA-seq methods (Baran-Gale et al., 2018).

### Data integration

In order to consider the relative differences between datasets, we applied a data integration technique and integrated all datasets together. The workflow of data integration was based on Seurat v3 (Butler et al., 2018 & Stuart et al., 2019). Firstly, this method aims to identify ‘anchors’ between each pair of datasets. This process reflects pairwise correspondences between individual cells that are hypothesized to originate from the same biological state. These ‘anchors’ are then used to reconcile the datasets, or transfer information from one dataset to another. We performed all the necessary steps of the data integration workflow:

1. The first step was a preprocessing of the datasets that was to create a list with gene expression matrices from each dataset and separately run Seurat preparation procedure — normalization using the NormalizeData function and feature selection with FindVariableFeatures.
2. We identified the anchors using the FindIntegrationAnchors function, which takes a list of Seurat objects as input. We mostly used default parameters for identifying anchors, except for ‘anchor.features’=6000 and ‘k.filter’=40.
3. We passed these anchors to the IntegrateData function, returning a Seurat object, which holds an integrated (or ‘batch-corrected’) expression matrix for all cells, enabling them to be jointly analyzed.
4. We analyzed this integrated dataset with the sc-RNA-seq pipeline described below.

### Pipeline for analyzing data

#### Filtering

In preparation for the main part of analyzing single cell gene expression data, it was important to make sure that all count data represent viable cells. Therefore, we considered the following quality control metrics:

Firstly, we plotted a histogram of the distribution of the number of genes per cell. The distribution was inspected for outlier peaks that were filtered out by thresholding afterwards. Cells with few detected genes could possibly be the sign of broken membrane, dead cells, etc.

Secondly, we filtered out genes that were not expressed in more than 10 cells.

#### Normalization

Most of the research, starting with the normalization part, was done using the R package Seurat, it showed good performance in analyzing single cell RNA-seq data and has most of the necessary functions with relevant methods: normalization, feature selection, clustering and data integration.

Since the sequencing data from the Tabula Muris dataset was generated with Smart-seq2 protocol, which does not allow for usage of UMI’s, we used the well-proven global scaling normalization method “LogNormalize” that normalizes the gene expression measurements for each cell by the total expression, multiplies this by a scale factor (10,000 by default), and log-transforms the result.

For the embryonic datasets, which use the UMI-based sc-RNA-seq protocol, we applied a recent normalization approach called ScTransform (Hafemeister et al., 2019). This method is available as part of the R package sctransform and has a direct interface to Seurat with function SCTransform. The algorithm ignores usage of scaling factors and concentrates on the construction of a generalized linear model relating sequencing depth of the cells to gene counts. After it calculates the Pearson residuals of the model, that represents transformation stabilized for variance.

#### Restoring cell trajectories with PAGA

In an analysis of single cell expression datasets that contain cell populations with transient processes (like EHT) it is important to capture this conversion and reflect it in the downstream analysis. In our research we performed trajectory inference with the Python library Scanpy and its framework PAGA (Wolf et al., 2018 & Wolf et al., 2019). This method provides an interpretable graph-like map of the arising data manifold, based on estimating connectivity of manifold partitions. We ran the PAGA with standard workflow:

1. Compute the nearest neighbors’ graph with pp.neighbors method.
2. Cluster the single-cell graph using the method tl.leiden. Optimal resolution was chosen after brute force of silhouette score of the clustering.
3. Construct the abstracted graph based on the connectivity between the clusters with tl.paga method, with threshold 0.01.
4. Compute UMAP-embeddings with use of the method tl.umap using PAGA-graph as initialization of UMAP (McInnes et al., 2018).

Afterwards, we incorporated these PAGA-initialized single-cell embeddings to our pipeline.

#### Clustering

With the Tabula Muris dataset we used the annotation, provided by the authors of the original article.

With the Embryo_dataset1, we used the SNN-clustering method available in Seurat:

1. First step is calculating the PCAs of the expression matrix with the function RunPCA.
2. Second step is running the function FindNeighbours that computes SNN-graph.
3. Finally, to cluster the cells we applied the function FindClusters to the calculated graph with use of Leiden algorithm and resolution 0.3 (Traag et al., 2019).

#### Dimensionality reduction

With all datasets we used UMAP-embeddings obtained from **PAGA**’s trajectory inference.

#### Marker genes identification

For the identification of marker genes of the clusters in each case we used the Seurat function *FindAllMarkers* with use of *MAST* algorithm (Finak et al., 2015). This technique was shown to perform better specifically on the single cell expression data. Afterwards, the heatmaps of single cell marker gene expression were plotted for each dataset with the function *DoHeatmap*.

#### Co-expression analysis

To understand the expression patterns of the seed transcription factors, we calculated the co-expression of TFs in the endothelial population in each dataset, summarizing the expression of TFs in every cell.

#### Inferring regulatory networks using ScTarNet

The important part of our research was the comprehension of how the seed transcription factors interact with the target genes. To answer this question, we inferred gene regulatory networks with the R package ScTarNet (Bergiers et al., 2018). This package provides a method that builds networks based on an input set of transcription factors and tests for indirect relationships and TF-TF interactions using distance correlations.

We applied a standard workflow of the ScTarNet:

- First step was initial correlation network inferring with function calculateTFstoTargets, which estimates relationships between TFs and target genes
- Second step is identifying TF-TF interactions using partial distance correlation with function calculateConditionalCors
  1. This distance correlation metric is a new statistical approach which enables detection of both linear and non-linear dependencies between variables that is particularly important for analyzing highly-dimensional data like single cell expression. Its main limitation is that it is an O(n2) operation, where n is the number of samples. We optimized the ScTarNet running time with replacing the distance correlation function to the one from the package Rfast (https://github.com/RfastOfficial/Rfast).

The output from this package are the target genes corresponding to the seed genes with three possible types of relationship: positive, negative and none.

#### Identifying the cells expressing TF-dependent genes

One of the questions of the research was to identify the cells with specific gene signatures of the groups of target genes associated to TFs. We solved this problem with the use of the R package **AUCell** (Aibar et al., 2017). This package uses the AUC-metric to determine whether a crucial subgroup of the input gene set is enriched within the expressed genes for each cell. The distribution of AUC-scores between every cell enables exploration of the relative expression. This scoring method is based on rankings which are built on the expression matrix. The outcome is the groups of cells linked to seed TFs. To find this groups we used default workflow of the package:

1. For each cell, the genes were ranked from highest to lowest value. The genes with the same expression value were shuffled. Therefore, genes with expression ‘0’ were randomly sorted at the end of the ranking. This step was done using the function *AUCell_buildRankings*.
2. To determine whether the gene set is enriched at the top of the gene-ranking for each cell, AUCell uses the “Area Under the Curve” (AUC) of the recovery curve. The function *AUCell_calcAUC* calculates this score, and returns a matrix with an AUC score for each gene-set in each cell.
3. The AUC represents the proportion of expressed genes in the signature, and their relative expression value compared to the other genes within the cell. We used this property to explore the population of cells that are present in the dataset according to the expression of the gene-set with function *AUCell_exploreThresholds*. The ideal situation is a bi-modal distribution, in which most cells in the dataset have a low AUC compared to a population of cells with a clearly higher value. Consequently, we picked thresholds that separated these two modes.
4. Cells with at least that value of threshold were selected for the further analysis.

Since **AUCell** is based on calculation of AUC it does not tell anything about the significance of the outcome. Therefore, we additionally performed Kruskal-Wallis tests over ranking matrix versus target genes. The null hypothesis was that the mean ranks of groups are the same. Afterwards, p-values retrieved from the tests were corrected for multiple testing with Holm-Bonferroni correction:

1. All p-values are sorted in order of smallest to largest. *m* is the number p-values.
2. If the 1-st p-value is greater than or equal to *α*/*m*, the procedure is stopped and no p-values are significant. Otherwise, go on.
3. The 1st p-value is declared significant and now the second p-value is compared to *α*/(*m* − 1). If the 2nd p-value is greater than or equal to *α*/(*m* − 1), the procedure is stopped and no further p-values are significant. Otherwise, go on. Cells that were found to be significantly and specifically expressing groups of target genes associated with seed TFs, were highlighted on the UMAP-embeddings of the dataset.

#### Detecting Runx1-specific signatures

To uncover any Runx1-specific patterns of single cell expression we applied straightforward approach:

1. All endothelial cells expressing Runx1 were allocated to a separate cluster (“x cells”).
2. Next, we performed **Seurat** marker gene identification contraposing cluster of endothelial cells to the “x cells” cluster.
3. Obtained gene lists were compared between adult tissues and Mouse Embryo dataset, calculating the number of overlaps.

#### Calculating the score of the tissues

To rank the tissues in term of reprogramming potential, we took scores of expression frequencies, co-expression of the seed TFs and presence of putative targets of GRNs. Afterwards, we normalized these scores to maximum and ranked.

## Code availability

All the code used for this study is available at the following GitHub repository: https://github.com/adamov-artem/EHT-study

## Acknowledgements

We thank Nicolas Descostes (EMBL Rome Bioinformatics service, Italy) for advice and technical support. Figures were created using BioRender.com.

## Conflict of interests statement

The authors declare no competing interests.

## Contributions

Artem Adamov, Conceptualization, Formal analysis, Visualization, Writing—review and editing; Yasmin Natalia Serina Secanechia, Conceptualization, Writing—review and editing; Christophe Lancrin, Conceptualization, Formal analysis, Supervision, Investigation, Visualization, Writing—original draft, Project administration, Writing—review and editing.

## Notes

### Competing Interest Statement

The authors have declared no competing interest.

