## Supplementary figures and images for "Single-cell transcriptome analysis of embryonic and adult endothelial cells allows to rank the hemogenic potential of post-natal endothelium"

### SupplementaryFigure1

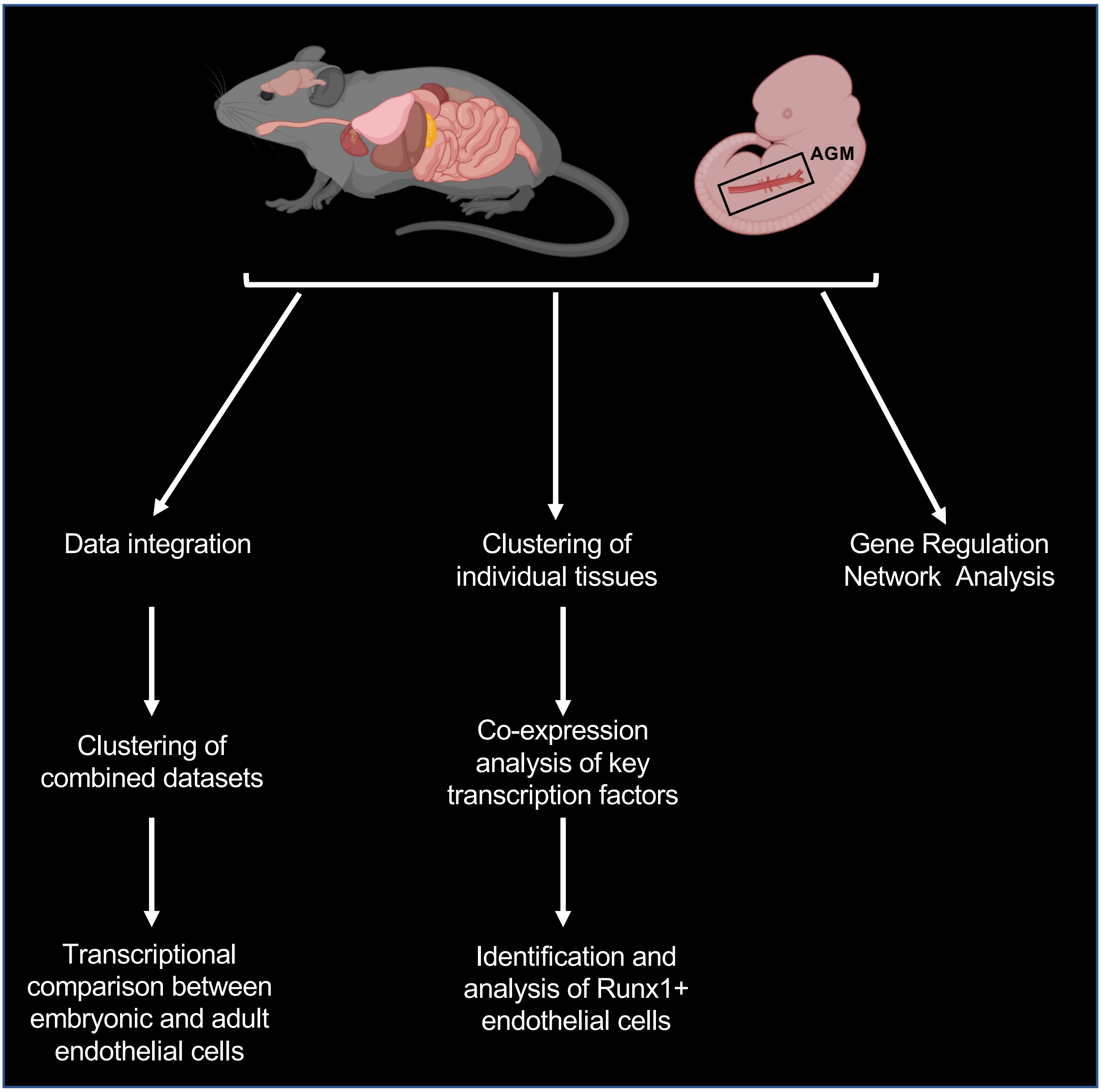


**Supplementary Fig. 1: Comparative analysis of murine embryonic and adult endothelial cells**
