## SupplementaryFigure2 for "Single-cell transcriptome analysis of embryonic and adult endothelial cells allows to rank the hemogenic potential of post-natal endothelium"

**a**

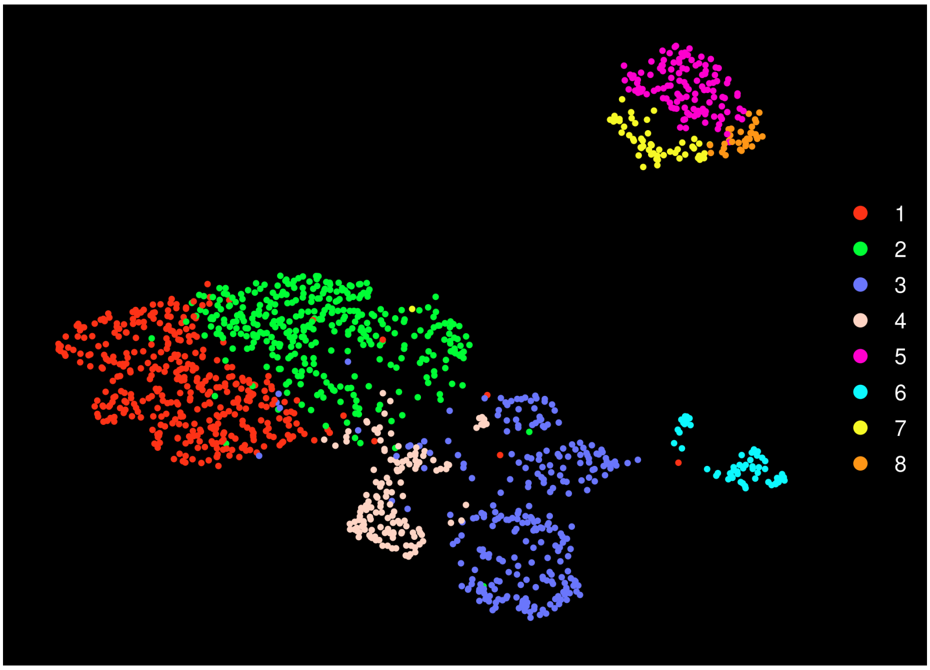


**b**

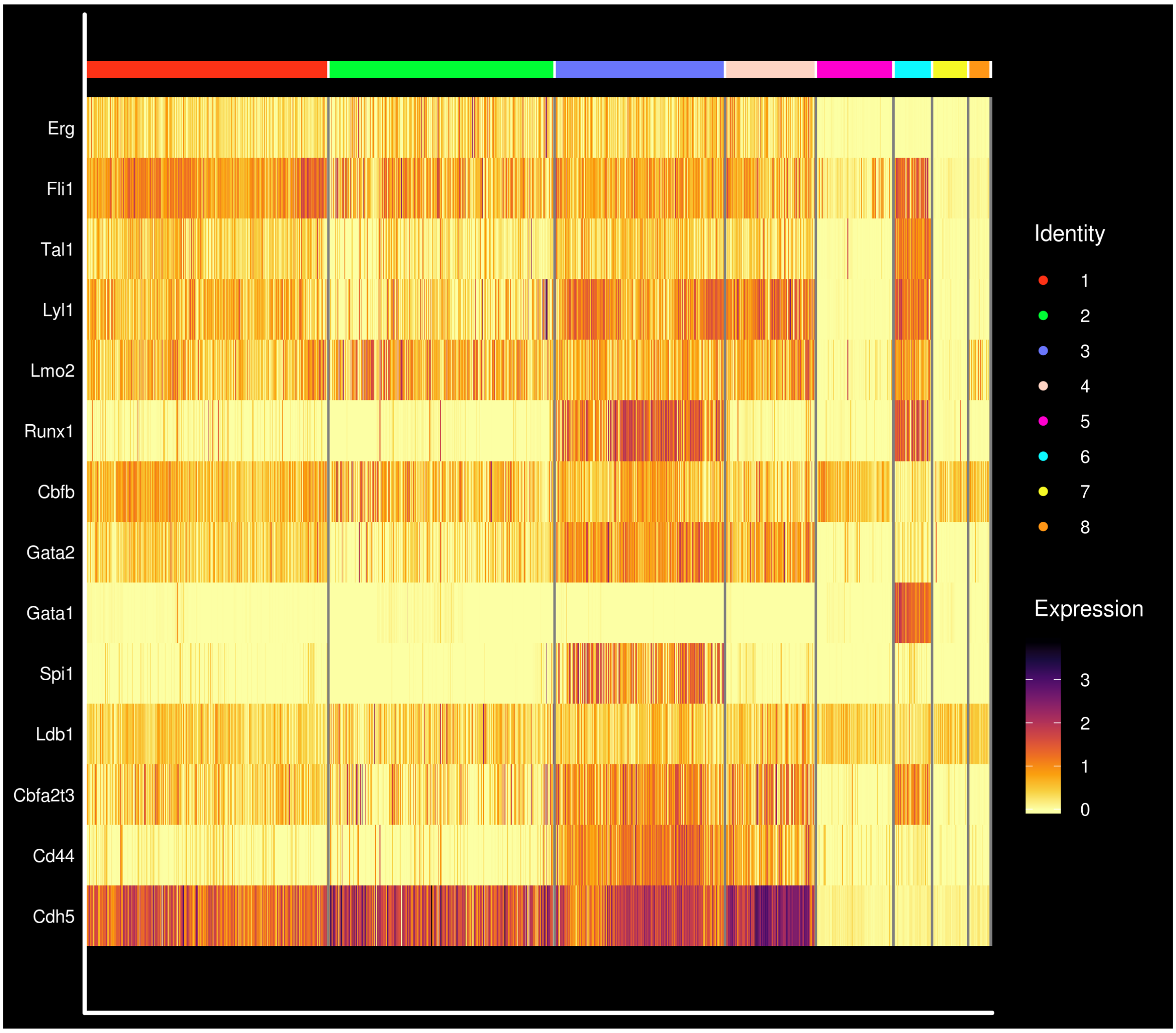


**Supplementary Figure 2: Clustering analysis of Embryo_dataset_1. a)** UMAP plots showing the clustering analysis result; **b)** Gene expression heatmap for a selection of key genes involved in the EHT process.
