## SupplementaryFigure3 for "Single-cell transcriptome analysis of embryonic and adult endothelial cells allows to rank the hemogenic potential of post-natal endothelium"

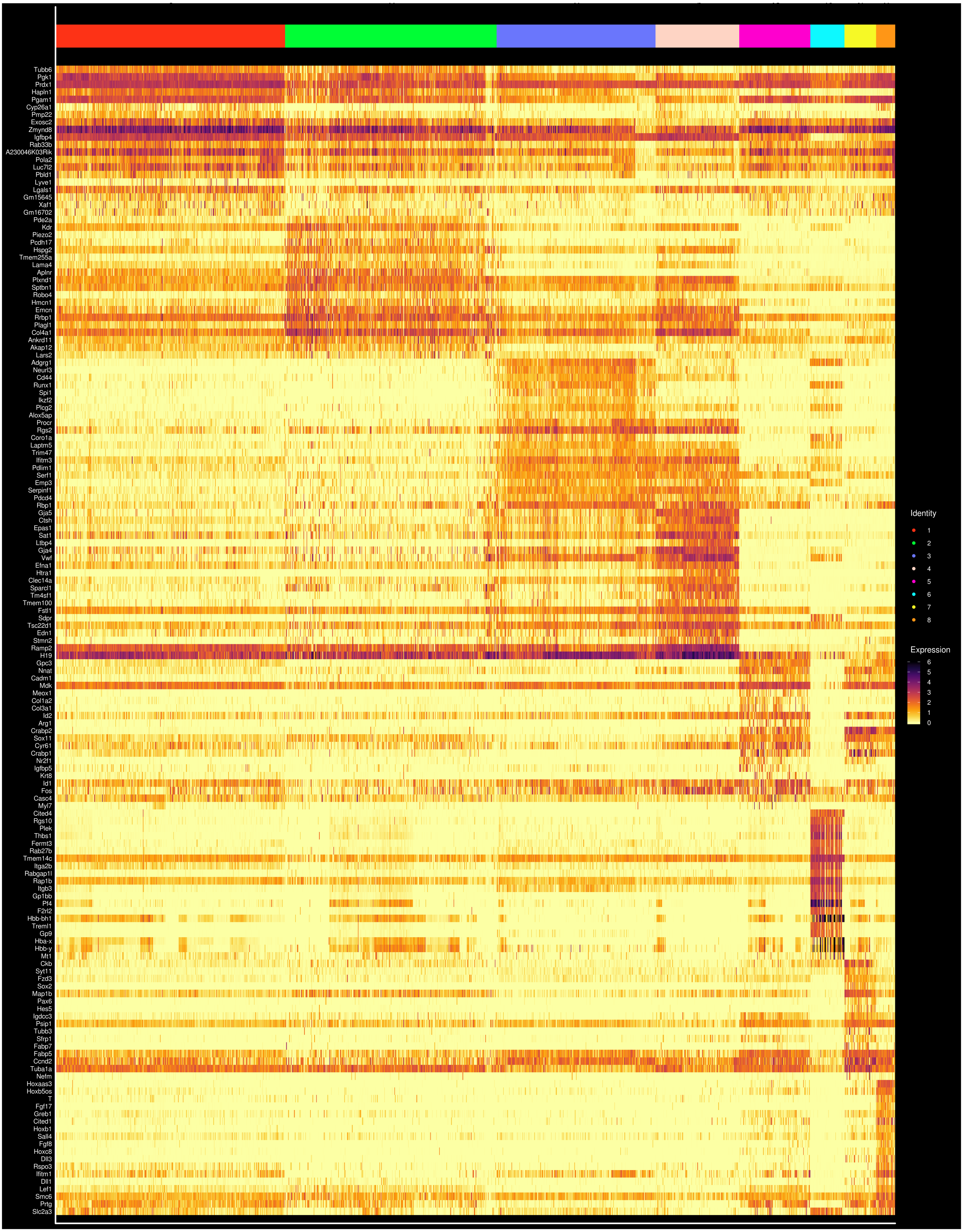


**Supplementary Figure 3: Marker genes of Embryo_dataset_1 clusters.** Expression heatmap for marker genes for the eight indicated clusters.
