## SupplementaryFigure4 for "Single-cell transcriptome analysis of embryonic and adult endothelial cells allows to rank the hemogenic potential of post-natal endothelium"

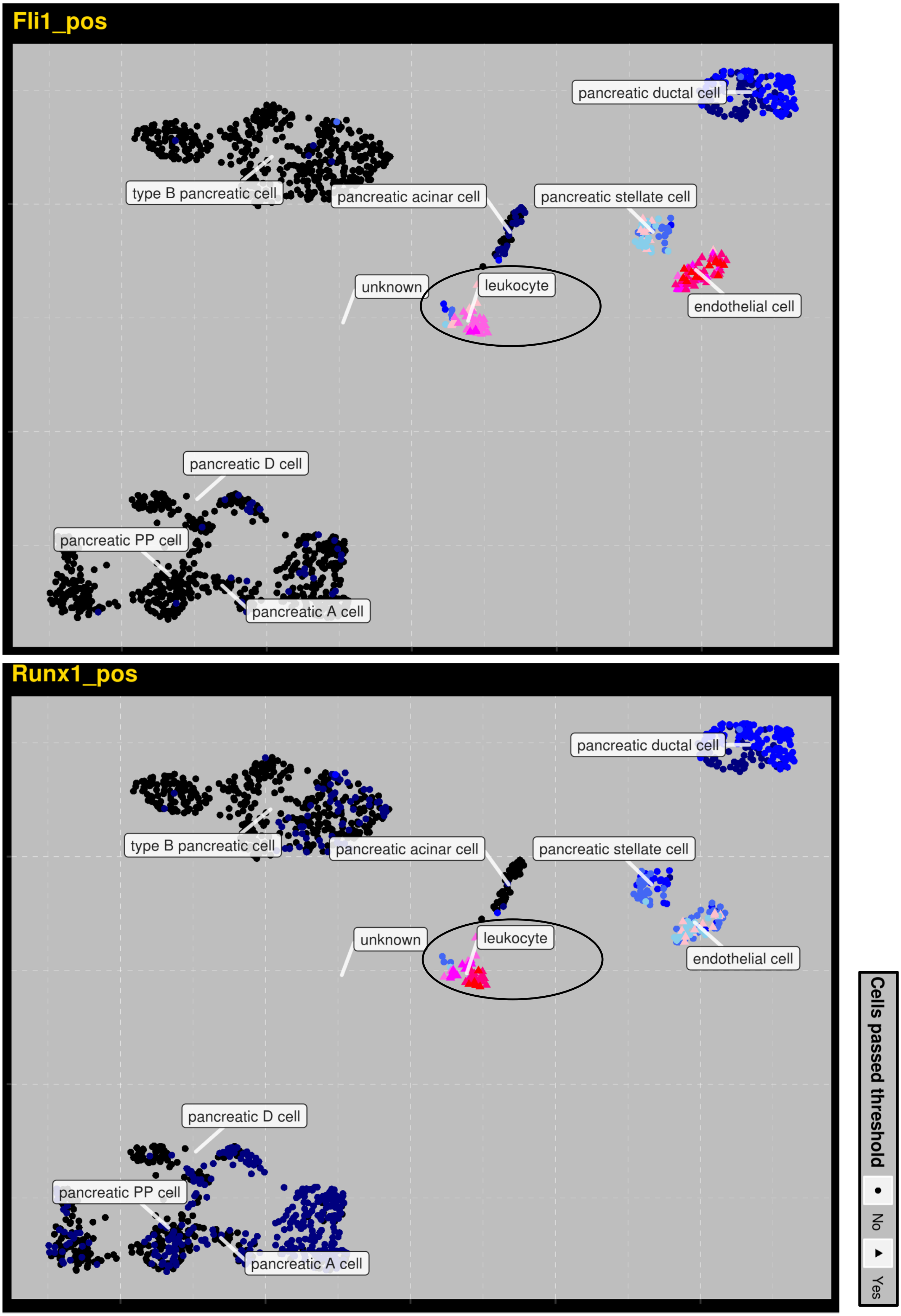


**Supplementary Figure 4: Overlap of genes between Fli1 and Runx1 only in leukocytes cluster.** UMAP plot highlighting the clusters in which the target genes of a given seed TFs are expressed. Only positive relationships are highlighted. The ellipse shows the leukocytes cell cluster. Colors represent AUC value – cells that passed AUC threshold colored in pink, others in blue.
