## SupplementaryTable1 for "Single-cell transcriptome analysis of embryonic and adult endothelial cells allows to rank the hemogenic potential of post-natal endothelium"

| Gene symbol | Number of common tissues | Name of tissues |
|---|---|---|
| *Cd63* | 5 | fat, kidney, limb_muscle, lung, trachea |
| *Gsn* | 5 | fat, heart, kidney, limb_muscle, trachea |
| *Mt1* | 5 | diaphragm, fat, kidney, limb_muscle, trachea |
| *Myl9* | 5 | brain, diaphragm, fat, limb_muscle, trachea |
| *Rgs5* | 5 | aorta, brain, diaphragm, limb_muscle, trachea |
| *Acta2* | 4 | diaphragm, fat, limb_muscle, trachea |
| *Art3* | 4 | brain, diaphragm, kidney, mammary_gland |
| *Bgn* | 4 | brain, kidney, limb_muscle, trachea |
| *Cd44* | 4 | diaphragm, kidney, limb_muscle, mammary_gland |
| *Ndrg2* | 4 | aorta, diaphragm, limb_muscle, trachea |
| *Notch2* | 4 | aorta, diaphragm, kidney, limb_muscle |
| *Pex7* | 4 | aorta, kidney, limb_muscle, lung |
| *Stip1* | 4 | aorta, fat, kidney, mammary_gland |

**Supplementary Table 1: Identification of marker genes in Runx1$^{+}$ adult endothelial populations**
